# Hippocampal single-cell RNA Atlas of chronic methamphetamine abuse-induced cognitive decline in mice

**DOI:** 10.1101/2025.05.02.651823

**Authors:** H Qiu, X Yue, YB Huang, ZL Meng, JH Wang, DF Qiao

## Abstract

**Background:** Chronic methamphetamine abuse leads to cognitive decline, posing a significant threat to human health and contributing to loss of productivity. However, the intricate and multifaceted mechanisms underlying methamphetamine-induced neurotoxicity have impeded the development of effective therapeutic interventions.

**Methods:** To establish a mouse model of cognitive decline induced by chronic methamphetamine exposure, we employed a large sample size and conducted two behavioral tests (Y-maze and novel object recognition test) at 2 and 4 weeks post-exposure. Subsequently, single-cell RNA sequencing was utilized to delineate the mRNA expression profiles of individual cells within the hippocampus.

Comprehensive bioinformatics analyses, including cell clustering and identification, differential gene expression analysis, cellular communication analysis, pseudotemporal trajectory analysis, and transcription factor regulation analysis, were performed to elucidate the cellular-level changes in mRNA profiles caused by chronic methamphetamine exposure.

**Results:** Our findings demonstrated impairments in working memory, spatial cognition, learning, and cognitive memory. After 4 weeks of behavioral testing, we identified diverse cell types in the hippocampi of METH- and saline-treated mice through scRNA-seq, including glial cells, stromal cells, vascular cells, and immune cells. We observed that methamphetamine exerts cell-specific effects on gene expression changes associated with neuroinflammation, blood-brain barrier disruption, neuronal support dysfunction, and immune dysregulation. Furthermore, cross-talk analysis revealed extensive alterations in cellular communication patterns and signal changes within the hippocampal microenvironment induced by methamphetamine exposure. Pseudotime analysis predicted hippocampal neurogenesis disorders and identified key regulatory genes implicated in chronic methamphetamine abuse.

Transcription factor analysis uncovered regulators and pathways linked to astrocyte-mediated neuroinflammation, endothelial junction integrity, microglial synaptic remodeling, and oligodendrocyte-supported neuronal cell bodies and axons. Additionally, it highlighted the role of neural precursor cells in various forms of neurodegeneration.

**Conclusions:** This study establishes a robust mouse model of cognitive impairment induced by chronic methamphetamine exposure. It provides valuable biological insights, characterizes the single-cell atlas of the hippocampus, and offers novel directions for investigating neurological damage associated with chronic methamphetamine-induced cognitive decline.

## Introduction

Methamphetamine (METH), a widely abused psychostimulant globally, particularly in southeast and east Asia, currently holds the position of being the world’s foremost illicitly manufactured synthetic drug. Its detrimental impact on human health, social stability, and economic development is immense(1). METH induces intense euphoria and addiction, while causing significant harm to various tissues and organs, especially the central nervous system. With advancements in productivity and medical standards, research focusing on neuropsychiatric issues associated with chronic METH abuse has become a prominent topic concerning drug users’ treatment, recovery, and quality of life. Chronic and regular usage of METH has been proven to result in an array of brain structural abnormalities such as cortical and hippocampal atrophy, nucleus accumbens hypertrophy, as well as decreased gray matter(2–4), leading to neuropsychiatric disorders including neurodegeneration, cognitive deficits, agitation, paranoia, depression, delusions, and schizophrenia. However, due to the wide range of affected targets and complex mechanisms underlying METH’s effects, the evolution and interplay of these effects during chronic processes make it challenging to analyze the etiology of these neuropsychiatric diseases.

Extensive clinical and experimental evidences suggest that chronic METH abuse leads to cognitive deficits(5–7), which share similarities with neurodegenerative disorders such as impaired inhibitory control, learning and memory decline, attention decrease, increased pathological protein levels, neurotoxicity, neuroinflammation, and blood-brain barrier (BBB) injury. The hippocampus plays a crucial role in regulating cognitive function as part of the limbic system. Previous studies have demonstrated that METH exerts complex negative effects on the hippocampus beyond its direct impact on dopaminergic and glutamatergic systems(8). It also indirectly affects various cell types within the hippocampus and alters the microenvironment. However, due to the intricate composition of the hippocampus and diverse mechanisms underlying METH’s effects, certain impacts on vulnerable cell types and critical signaling pathways may be concealed during chronic exposure, making it challenging to unravel these mechanisms. Currently, there is limited understanding of the specific mechanisms responsible for chronic METH-induced neurotoxicity in the hippocampus leading to cognitive deficits; furthermore, theories explaining this wide range of effects are scarce.

In this study, we employed single-cell RNA sequencing (scRNA-seq) to construct a comprehensive hippocampal atlas of chronic METH abuse-induced cognitive decline in mice. To elucidate the underlying mechanisms by which chronic METH abuse affects the hippocampus and leads to cognitive deficits, we conducted animal behavior tests, performed scRNA-seq analysis with clustering and differential gene expression analyses, investigated cellular cross-talk, examined single-cell trajectories, and analyzed transcription factors. This study provides in vivo evidence of heterogeneous changes in hippocampal cells under chronic METH abuse and contributes to a more robust understanding of the mechanistic basis for cognitive deficits while also bridging previous theories on METH neurotoxicity.

## Materials and methods

### Animal and Treatments

All mice used were male and on C57BL/6 background. Healthy 7-week-old mice were purchased from the Laboratory Animal Center of Southern Medical University (Guangzhou, China). Mice were quarantined for 2 weeks and then housed in the Central Exhaust Ventilation Cage System (Houhuang experimental equipment technology, Suzhou, China) with 12 hours (h) light/12 h dark cycle in a temperature-(20–22 ℃) and humidity- (45–55%) controlled vivarium and had ad libitum access to water and food. All protocols approved by the Institutional Animal Care and Use Committee in Southern Medical University (Ethical number: L2022125), and consistent with NIH Guidelines for the Care and Use of Laboratory Animals (8th Edition, U.S. National Research Council, 2011). After 1 week for adapting, mice were 10 weeks of age at the start of the experiment. They were randomly divided into two groups, and then treated by saline or METH (National Institutes for the Control of Pharmaceutical and Biological Products, Beijing, China). Mice of METH group were intraperitoneally administered METH dissolved in saline at a incremental dose from 1 up to 10 mg/kg, and the daily dose was evenly divided into two doses at 12-hour intervals, while those of saline control underwent the same way of injection with saline simultaneously. The treatments lasted for 28 days. However the last administration of 12 h before animal behavior tests or sample-collecting would be skipped. For scRNA-seq, we used 20 mice (10 from saline group, 10 from METH group, and each random 2 mice from same cage were composed of 1 sample).

### Behavioral Experiments

Behavioral testing of mice used for scRNA-seq experiments followed an established protocol. To minimize any stimulation, mice would be handled for 5 days to be habituated to environment and tester (30 min to environment, 5 min to tester) before behavioral experiments. Less irritating behavioral experiments were used to evaluate in order to minimize impacts on scRNA-seq.

### Y-Maze

Y maze test was used to evaluate spontaneous alternation behavior in mice. The Y-maze was made of three arms set at angles of 120°. Its walls are covered with frosted adhesive paper. Mice were placed in the end of fixed arm of Y-maze, then they were allowed to freely explore whole Y-maze for 8-min session while camera system recorded the behavior of mice. The total number of entries of each arm (recognizing that all four legs of mouse entered the arm as the standard for entries), the number of alternations (an alternation is defined as successive entries into the three arms on overlapping triplet sets.), and the ratio of them were regarded as statistical indicators to assess spatial working memory.

### Novel Object Recognition Test

Novel object recognition test (NOR) was applied to assess mice recognition memory. The testing room was lit with four LED lights which provide the light intensity of approximately 20 lx in each test box (50 × 50 × 40 cm). Mice were put into test room to habituate for 24h before beginning of test, and would be given 10 min to explore the object-free box and were then returned to their home cage. In training stage, two objects of same appearance were placed at a symmetrical position in test box where mice would explore freely for 10min to be fully acquainted with objects. After training, mice were sent back to their home cage to rest for 30 min. Then in testing stage, they were placed back into original test box with a familiar old object and a novel one at the same place and also allowed to explore freely for 10 min. The familiar object and novel object and their placement were counterbalanced within each group. Camera system was placed above the test box and recorded behaviors of mice in test box for statistics.

The following behaviors were scored as exploration: biting, sniffing, licking, touching the object with the nose or with the front legs, or proximity (≤1 cm) between the nose and the object. If the mouse put its hind legs on objects, or stood on top of the object or completely immobile, exploration was not scored. The preference index for the novel object was calculated as (time spent exploring the new object / the total time spent exploring both objects), and the discrimination index was calculated as (time spent exploring the new object − time spent exploring the familiar old object) / (total time spent exploring both objects). Behavior was scored on video by two observers blinded to the mice’s treatment.

### Brain Samples Collecting

For scRNA-seq, we used hippocampus tissues of 20 mice (10 from saline group, 10 from METH group, and each random 2 mice from same cage were regarded as 1 sample). Hippocampus collecting was carried out under the guidance of JOVE(8).

After the mice were subjected to 3 min of pre-cooling with PBS for cardiovascular perfusion, whole hippocampus was dissected integrally on icein the shortest time (within 5 min) and preserved in tissue preservation solution (SeekGene, China) at 4℃ for following tissue dissociation.

### Cell Preparation for Sing Cell RNA Sequencing

After harvested, tissues were washed in ice-cold RPMI1640 and dissociated using Tissue Dissociation Reagent A (Seekone K01301-30, China) from SeekGene as instructions. DNase Ⅰ (Sigma 9003-98-9, USA) treatment was optional according to the viscosity of the homogenate. Cell count and viability was estimated using fluorescence Cell Analyzer (Countstar® Rigel S2) with AO/PI reagent after removal erythrocytes (Solarbio R1010, China) and then debris and dead cells removal was decided to be performed or not (Miltenyi Biotec 130-109-398/130-090-101, USA). Finally fresh cells were washed twice in the RPMI1640 and then resuspended at 1×10^6^ cells/mL in 1×PBS and 0.04% bovine serum albumin.

### Single Cell RNA Sequencing Library Construction and Sequencing

Single-cell RNA-Seq libraries were prepared using Chromium Next GEM Single Cell 3ʹ Reagent Kits v3.1 (10x Genomics Catalog No.1000268, USA). Briefly, appropriate number of cells were mixed with reverse transcription reagent and then loaded to the sample well in Chromium Next GEM Chip G. Subsequently Gel Beads and Partitioning Oil were dispensed into corresponding wells separately in chip. After emulsion droplet generation reverse transcription were performed at 53℃for 45 min and inactivated at 85℃ for 5 min. Next, cDNA was purified from broken droplet and amplified in PCR reaction. The amplified cDNA product was then cleaned, fragmented, end repaired, A-tailed and ligated to sequencing adaptor. Finally, the indexed PCR was performed to amplify the DNA representing 3’ polyA part of expressing genes which also contained Cell Bar code and Unique Molecular Index. The indexed sequencing libraries were cleanup with SPRI beads, quantified by quantitative PCR (KAPA Biosystems KK4824) and then sequenced on illumina NovaSeq 6000 with PE150 read length.

### Processing the Single Cell RNA Sequencing Data and Date Quality Control

The raw sequencing data was processed by Fastp firstly (Chen, Zhou et al. 2018) to trim primer sequence and low quality bases. And then we employed the Cell Ranger software v7.0 (10X Genomics) obtained from https://www.10xgenomics.com/support/software/cell-ranger/downloads to precess sequence data and aligned to mouse mm10 genome in order to obtain gene expression matrix. Subsequently, the gene expression matrices of each sample were obtained and further analyzed using Seurat v.4.4.0. Then we determined effective cells using the following criteria: (a) total UMI counts more than 1000 but less than the 97th percentile for each cell; (b) gene numbers between 1000 and 7000; (c) percentage of mitochondrial genes < 10%; and (d) percentage of hemoglobin genes < 0.1%. Moreover, we excluded genes expressed in fewer than 3 cells, as well as MALAT1, mitochondrial genes, and hemoglobin genes. And the remaining cells and genes were used for subsequent analysis. Quality control charts are included in SI.4.

### Data integration, Dimensionality reduction, and Clustering

After performing quality control and filtering, DoubletFinder v2.0.3 was then employed to identify double cells with an expectation value of doublets calculated as an increase of 0.8% for every 1000 additional cells, pN at 0.25, and pK according to function Find.pk. Library-size normalization was conducted for each cell using the NormalizeData function in Seurat v.4.4.0. The 2000 most variable genes were identified using the FindVariableGenes. Subsequently, all libraries were integrated using Harmony (Ilya Korsunsky et al., 2019) to correct batch effects. ScaleData was applied to regress out variability associated with the numbers of UMIs, followed by dimensionality reduction through RunPCA and RunUMAP (dimensions = 1:20). Finally, clustering of cells was performed using the FindClusters function with a resolution of 0.8 based on the 1:24 dimensions.

### Celltype Identifcation

FindAllMarkers was used to compare each cluster to all others to identify canonical cell type-specific marker genes. Each retained marker gene was expressed in a minimum of 25% of cells and at a minimum log fold change threshold of 0.25. The clustering differential expressed genes were considered significant if the adjusted P-value was less than 0.05 and the avg_log2FC was ≥ 0.25. And the clusters were annotated using cell type-specific signatures and marker genes from Cellmarker2.0 and PanglaoDB.

### Differentially Expressed Genes (DEGs) analysis and Enrichment Analysis

DEG analysis between cell types was performed following the muscat (v.1.16.0, Crowell et al. 2020) R package pipeline, which facilitates multi-sample, multi-condition comparisons of single-cell RNA-seq data. Genes were ranked by absolute log2 fold-change (log2FC), and those with p-values > 0.05 (adjusted for multiple comparisons) and log2FC < 0.25 were removed. Gene Ontology (GO, https://www.geneontology.org/) enrichment analysis of DEGs was implemented by the clusterProfiler R package (Yu et al., 2012). GO terms with corrected P value less than 0.05 were considered significantly enriched by DEGs. Kyoto Encyclopedia of Genes and Genomes database (KEGG, http://www.genome.jp/kegg/, Kanehisa et al., 2008) is a database resource for understanding high-level functions and utilities of the biological system, such as cell, organism and ecosystem, from molecular-level information, especially large-scale molecular datasets generated by genome sequencing and other high-through put experimental technologies. We used clusterProfiler R package to test the statistical enrichment of DEGs in KEGG pathways.

### Cellular Cross-talk Analysis

We utilized the Cellchat (v.1.6.1, Suoqin Jin et al., 2021, http://www.cellchat.org/) to analyze the expression abundance of ligand–receptor interactions across hippocampal cell types. The analysis was conducted in accordance with the CellChat user guidelines(9). Cell communication networks were inferred by identifying differentially expressed ligands and receptors between distinct hippocampal cell types. Interaction probabilities at the ligand–receptor level were calculated using the default ‘truncatedMean’ method, under which the average expression of a signaling gene is set to zero if it is expressed in less than 10% of cells within a given group. The influence of population size was normalized during the calculation of interaction probabilities. Furthermore, L-R interaction probabilities within each signaling pathway were aggregated to compute pathway-level communication probabilities using the function ‘computeCommunProbPathway’. Cell–cell communication networks were subsequently summarized by summing the number of interactions or the previously calculated communication probabilities via the function ‘aggregateNet’. To compare signaling patterns between METH and Saline groups, we conducted differential expression analysis between METH and Saline samples within each cell type. To ensure consistency in node size and edge weights across inferred networks from different datasets, we calculated the maximum number of cells per cell group and the maximum interaction counts (or interaction weights) across the two datasets. Differential and conserved networks were identified based on their functional roles, and their divergence indicators were computed accordingly. We also compared outgoing, incoming, and overall signaling patterns between the two datasets and across different cell types. To identify major signaling roles of specific cell types within defined pathways, we visualized the computed centrality scores using heatmaps. Discrepant ligands and receptors were identified if both the log-fold change (logFC) and the percentage of cells expressing the gene in a given cluster exceeded 0.1 in either sender or receiver cells, as determined by the function ‘identifyOverExpressedGenes’. Finally, differentially expressed L-R pairs were extracted based on upregulated or downregulated ligands and receptors in METH compared to Saline groups.

### Pseudotime analysis

Monocle2(10) (version 2.18.0) was utilized for pseudotime trajectory analysis to elucidate the differentiation trajectory of cell development. The UMI matrix was extracted from the Seurat object and used to create a new object with the ‘newCellDataSet’ function. For the trajectory analysis, the 2000 most highly variable genes were selected, followed by dimensionality reduction with the DDRTree method and cell ordering with the ‘orderCells’ function. Genes exhibiting significant changes along the pseudotime trajectory were identified using the ‘differentialGeneTest’ function. The ‘plot_cell_trajectory’ function was utilized to visualize the trajectory in a two-dimensional space, effectively presenting the cell differentiation state. The observed branching patterns in the single-cell trajectories reflect distinct biological functions, which were inferred based on gene expression dynamics during development.

### SCENIC analysis

SCENIC analysis was utilized to explore key transcription factors (TFs) that regulate inter-cell type differences. The analysis was conducted using pySCENIC with default parameters, following the established protocol(11). Initially, a co-expression network was constructed using GRNBoost2. Subsequently, regulons for each transcription factor were inferred using the RcisTarget motif databases(12) (mm10-refseq-r80-10kb-up-and-down-tss.mc9nr.feather and mm10-refseq-r80-500bp-up-and-100bp-down-tss.mc9nr.feather). Activity scores (AUC values) for each TF regulon in individual cells were calculated using AUCell. Significant TF regulons associated with specific cell types were identified using the ‘FindAllMarkers’ function in Seurat. The regulatory networks involving transcription factors and their target genes were further visualized and analyzed using Cytoscape (v3.10.1). Additionally, functional enrichment analyses of the target genes were conducted using the clusterProfiler R package, as previously described.

### Statistical Analysis

For the experimental data, statistical analyses and graphical visualization were performed using GraphPad Prism software (version 8.0.2, USA). The normality of data distribution was assessed using the Shapiro – Wilk test, although the sample size in each group was close to 30, which is considered relatively large. Results from the Y-maze and NOR tests were presented as mean ± SEM. Between-group differences in the alternation triplet of the Y-maze, as well as the discriminant index, preference index, total exploration time, and frequency of interactions with new and old objects within 10 minutes, were analyzed using the unpaired Student’s t-test. A P value < 0.05 was considered statistically significant, and all statistical tests were two-tailed.

For the single-cell RNA sequencing data, statistical analyses and figure generation were conducted using R (v4.3.1). Bioinformatic tools were applied to analyze cellular clustering, differentially expressed genes, functional enrichment, intercellular signaling cross-talk, single-cell trajectory inference, and SCENIC-based regulatory network analysis.

## Result

### 1. Chronic METH exposure led to cognitive decline in adult mice

The construction of the METH chronic exposure mouse model and the arrangement of behavioral experiments are illustrated in Fig.1.A. Adult male mice subjected to chronic METH exposure exhibited significant declines in working memory and spatial cognition, as demonstrated by Y-maze tests conducted at the end of both the second and fourth weeks of administration. In the 2-week test, the mean difference between the METH-treated (n=32) and saline-treated (n=29) groups was -0.1664 (95% CI: -0.2126 to -0.1202, p < 0.0001). In the 4-week test, the mean difference was -0.2202 (METH, n=30 vs. Saline, n=29; 95% CI: -0.2728 to -0.1676, p < 0.0001) (Fig.1B).

**Fig. 1.**
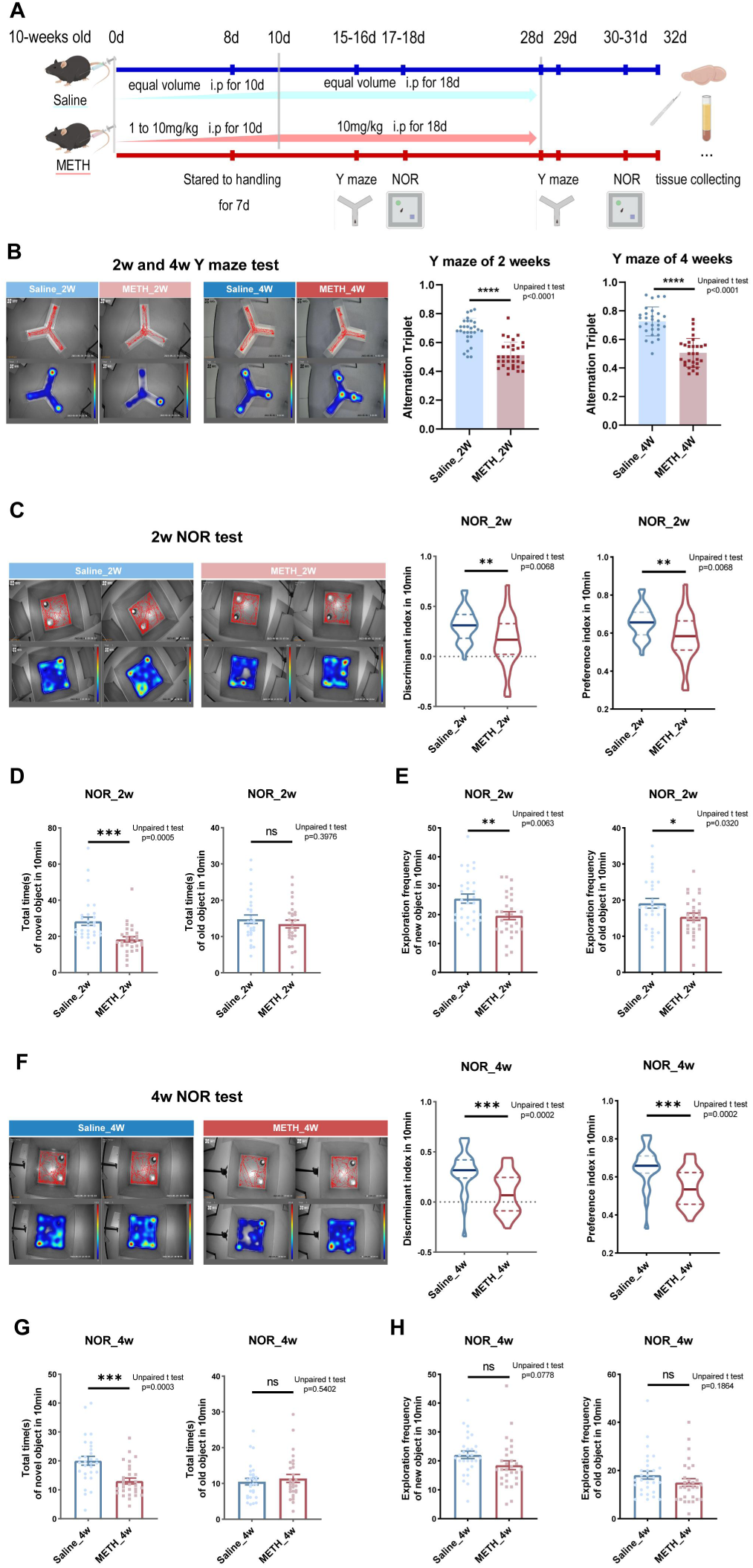
Mice treatment and result of behaviour tests. (A) The construction mouse model of chronic METH abuse. (B) The track visualizations and heatmaps of Y maze in 2 and 4 weeks. (C) The track visualizations, heatmaps and the discrimination and preference index of 2 weeks NOR test. (D-E) The total exploration time and frequency of familiar or novel objects of 2 weeks NOR test. (F) The track visualizations, heatmaps and the discrimination and preference index of 4 weeks NOR test. (G-H) The total exploration time and frequency of familiar or novel objects of 4 weeks NOR test. Asterisks indicate statistical significance (*p < 0.05; **p < 0.01; ***p < 0.001; ****p < 0.0001). All data are presented as mean ± SEM.

Following chronic METH exposure, NOR tests also revealed impairments in the animals’ ability to discriminate novel objects. In the 2-week NOR test, the discrimination index was 0.3119 ± 0.0300 (mean ± SEM) for saline-treated mice (n = 29), compared to 0.1569 ± 0.0462 for METH-treated mice (n = 29). The mean difference in discrimination index between the METH and Saline groups was -0.1550 ± 0.0551 (p = 0.0068, 95% CI: -0.2653 to -0.0446) (Fig.1C). The preference index was 0.6559 ± 0.0150 for saline-treated mice and 0.5785 ± 0.0231 for METH-treated mice, resulting in a mean difference of -0.07748 ± 0.0276 (p = 0.0068, 95% CI: -0.1327 to -0.0223) (Fig.1C).

The total exploration time for novel objects was 28.36 ± 2.285 seconds in the saline group, compared to 18.38 ± 1.453 seconds in the METH-treated group. The mean difference in total exploration time for novel objects was -9.974 ± 2.708 (p = 0.0005). For old objects, the total exploration time was 14.79 ± 1.173 seconds in the saline group and 13.44 ± 1.067 seconds in the METH group, yielding a mean difference of -1.352 ± 1.586 (p = 0.3976) (Fig.1D).

Exploration frequency analysis showed that saline-treated mice explored the novel object an average of 25.55 ± 1.565 times, compared to 19.66 ± 1.368 times in the METH group. The mean difference in novel object exploration frequency was -5.897 ± 2.079 (p = 0.0063) (Fig.1E). Similarly, the frequency of old object exploration was 19.14 ± 1.359 in the saline group versus 15.41 ± 1.010 in the METH group, with a mean difference of -3.724 ± 1.693 (p = 0.0320) (Fig.1E).

Collectively, these findings indicate that following 2 weeks of METH treatment, mice exhibited significant impairments in cognitive and mnemonic functions, as well as reduced exploratory behavior.

In the 4-week NOR test, the discrimination index was 0.3019 ± 0.0389 for the saline group (n = 29), compared to 0.0880 ± 0.0368 for the METH group (n = 29). The mean difference in discrimination index between the METH and saline groups was -0.2139 ± 0.0536 (p = 0.0002, 95% CI: -0.3213 to -0.1066) (Fig.1F). The preference index was 0.6510 ± 0.0195 in the saline group and 0.5440 ± 0.0184 in the METH group, resulting in a mean difference of -0.1070 ± 0.0268 (p = 0.0002, 95% CI: -0.1606 to -0.0533) (Fig.1F).

The total exploration time for novel objects was 20.04 ± 1.502 seconds in the saline group and 13.05 ± 1.019 seconds in the METH group, yielding a mean difference of -6.996 ± 1.815 (p = 0.0003) (Fig.1G). For old objects, the total exploration time was 10.50 ± 0.919 seconds in the saline group and 11.39 ± 1.119 seconds in the METH group, with a mean difference of 0.8922 ± 1.448 (p = 0.5402) (Fig.1G).

Exploration frequency for novel objects was 22.10 ± 1.293 in the saline group and 18.52 ± 1.521 in the METH group, resulting in a mean difference of -3.586 ± 1.996 (p = 0.0778) (Fig.1H). For old objects, the exploration frequency was 18.10 ± 1.658 in the saline group and 15.03 ± 1.585 in the METH group, with a mean difference of -3.069 ± 2.294 (p = 0.1864) (Fig.1H).

These findings indicate that chronic METH exposure leads to a decline in learning and cognitive memory abilities in mice. Although no significant differences in exploration frequency were observed between the METH and saline groups during the 4-week test, the discrimination and preference indexes showed greater statistical significance and further deterioration in the METH group compared to the 2-week test.

These results suggest that prolonged METH exposure may progressively impair cognitive function.

### 2. Identification of hippocampal cell clusters in mice with chronic METH exposure

To investigate the impact of chronic METH abuse, single-cell RNA-seq was performed on the hippocampi of mice treated with either saline or METH. A total of 60,549 high-quality cells were obtained after stringent quality filtering, including 31,219 cells from saline-treated mice and 29,330 cells from METH-treated mice. Clustering analysis was then conducted based on gene expression profiles using cluster-specific variable genes, following the Seurat pipeline (logfc.threshold = 0.25, min.pct = 0.25). Subsequently, Uniform Manifold Approximation and Projection (UMAP) analysis at a resolution of 0.8 identified a total of 31 distinct transcriptional clusters in each experimental group (Fig.2.A). Using established hippocampal markers from databases (primarily CellMarker2.0 and PanglaoDB), we annotated 18 distinct cell types within the hippocampus. These included astrocytes (cluster 6 and 27, marked by *Aqp4*, *Gfap*, *Gja1*, and *Slc1a3*), Cajal-Retzius cells (cluster 14, marked by *Cntnap2*, *Reln*, *Clstn2*, and *Kcnh7*), Cldn5^+^ and Cldn5^−^ mural cells (cluster 9 and 11, marked by *Vtn*, *Pdgfrb*, *Rgs5*, and *Atp13a5*, distinguished by high or low expression of *Cldn5*), endothelial cells (cluster 4, 7, 8, 10, and 16, marked by *Cldn5*, *Flt1*, *Pecam1*, and *Kdr*), ependymal cells (cluster 23, marked by *Ccdc153*, *Tmem212*, *Rarres2*, and *Mia*), fibroblasts (cluster 25, marked by *Dcn*, *Spp1*, *Nupr1*, and *Igfbp2*), macrophages (cluster 12, marked by *Pf4*, *Mrc1*, *F13a1*, and *Lyz2*), microglia (cluster 0, 1, 3, 21, and 29, marked by *Tmem119*, *Cx3cr1*, *P2ry12*, and *Aif1*), neurons or neuroblasts (cluster 17, marked by *Igfbpl1*, *Sox11*, *Fabp7*, and *Nnat*), neutrophils (cluster 26, marked by *S100a9*, *Retnlg*, *G0s2*, and *Slpi*), NK-T cells (cluster 19, marked by *Ccl5*, *Trbc2*, *Cd3g*, and *Ms4a4b*), neural stem cells (NSCs, cluster 20, marked by *Clu*, *Aldoc*, *Cpe*, and *Hopx*), neurons derived from NSCs (cluster 22, marked by *Chchd10*, *Pcp4*, *Fxyd1*, and *Ppp1r1b*), oligodendrocytes (cluster 2, 5, 24, 28, and 30, marked by *Mbp*, *Mog*, *Cldn11*, and *Mag*; cluster 24 was identified as perivascular oligodendrocytes), oligodendrocyte precursor cells (cluster 13, marked by *Pdgfra*, *Cacng4*, *Sox10*, and *Cspg4*), smooth muscle cells (cluster 15, marked by *Acta2*, *Myl9*, *Tagln*, and *Myh11*), and T cells (cluster 18, marked by *Cd44*, *Cd52*, *Cd74*, and *Plac8*). The top four highly expressed genes for each cluster are displayed (Fig.2.B&D, SI.1). These findings indicate that, in addition to neurons, the hippocampus comprises a diverse array of cell types, including neuroglia, stromal cells, vascular cells, and immune cells, forming a complex yet highly organized microenvironment. Within this microenvironment, METH exerts widespread effects across multiple cell types, potentially contributing to impairments in learning and memory. This knowledge may help elucidate the characteristics and functional alterations of specific hippocampal cell types under chronic METH exposure.

**Fig. 2.**
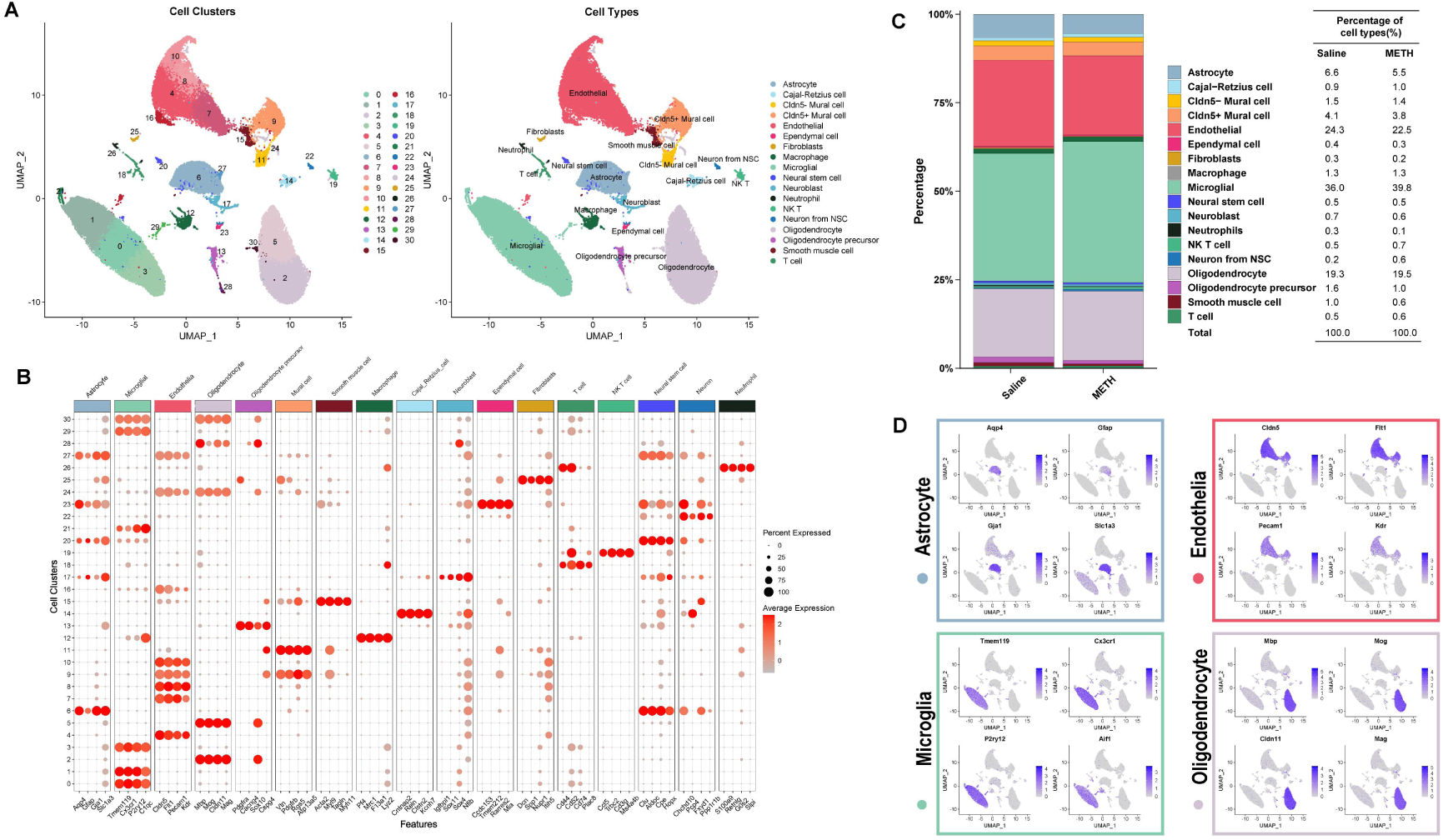
Hippocampus cell clusters of saline and METH mice. (A) The Umap-distributed plots showing 31 hippocampal cell clusters in saline and METH groups. (B) The average expression of known marker genes of hippocampal clusters. (C) The percentage of cells in hippocampal clusters in saline and METH groups. (D) The marker genes expression’s featureplots of astrocyte, endothelia, microglia and oligodendrocyte.

Following clustering analyses, cell proportion analysis revealed that chronic METH exposure led to a decrease in the proportion of astrocytes from 6.6% to 5.5%, endothelial cells from 24.3% to 22.5%, smooth muscle cells from 1.0% to 0.6%, oligodendrocyte precursor cells from 1.6% to 1.0%, neutrophils from 0.3% to 0.1%, ependymal cells from 0.4% to 0.3%, and fibroblasts from 0.3% to 0.2%. Conversely, an increase was observed in the proportion of microglia from 36.0% to 39.8%, NK-T cells from 0.5% to 0.7%, and neurons derived from NSCs from 0.2% to 0.6% (Fig. 2.C).

### 3. Chronic METH exposure induced gene and function changes of hippocampal cells in mice

Chronic METH abuse elicits a diverse array of cellular responses, ultimately resulting in multifactorial cognitive decline. To elucidate the underlying mechanisms of this cognitive impairment, we employed muscat — a pseudobulk-based analytical framework — for multi-sample, multi-condition, cluster-specific differential gene expression analysis, enabling the identification of differentially expressed genes (DEGs) within each cell cluster. Strikingly, most DEGs were found to be specific to individual cell types following chronic METH exposure. For example, Cldn5^+^ mural cells exhibited the highest number of DEGs, with 679 identified (368 upregulated and 311 downregulated) (Fig.3.A), including 336 cell type-specific DEGs (Fig.3.B).

**Fig. 3.**
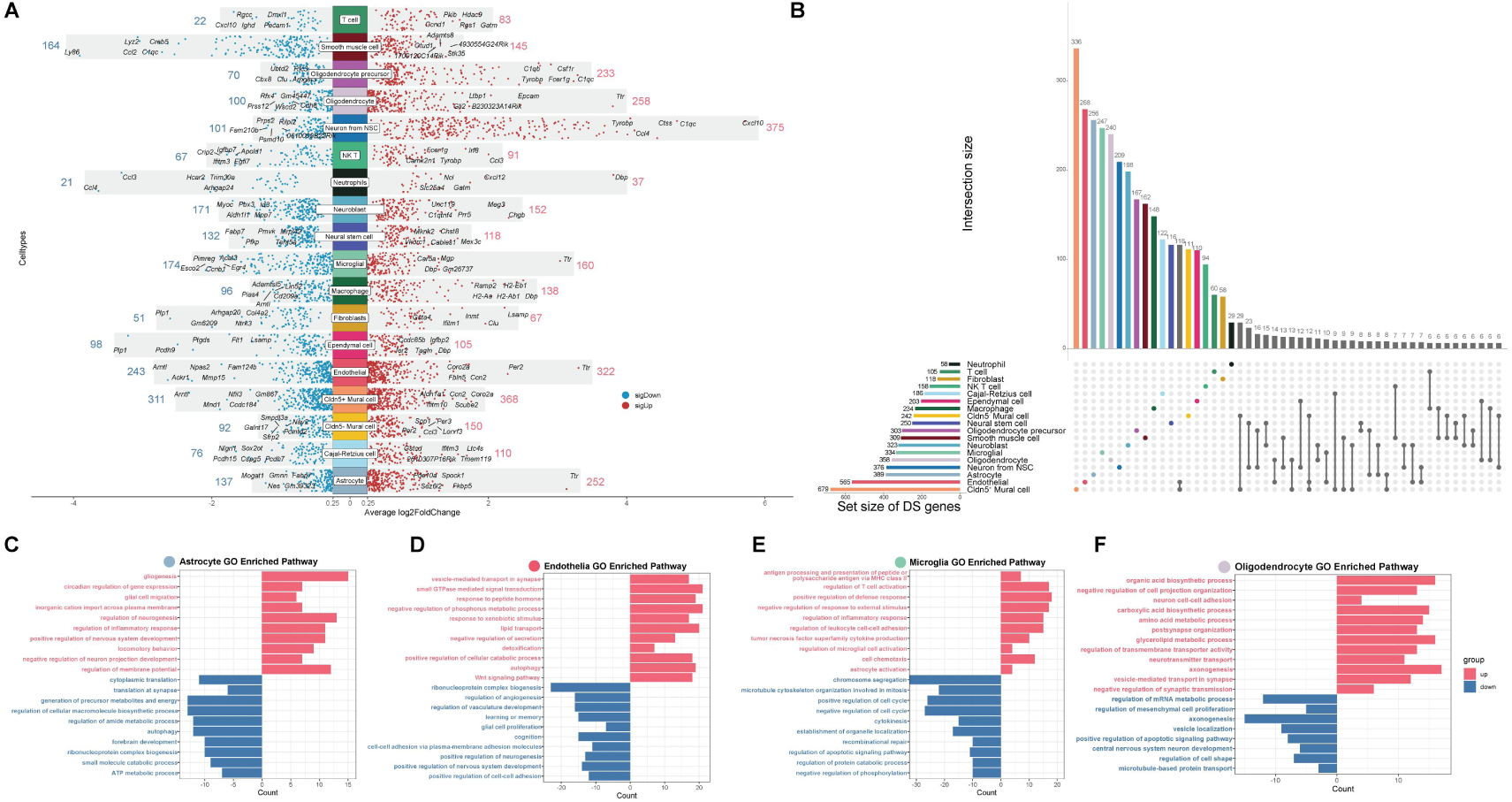
Differentally expressed genes(DEGs) induced by chronic METH exposure in hippocampal cell types. (A) The manhattanplot showing the number of DEGs in each hippocampal cell types(METH v.s. saline), the red dots and number represented for up-regulated genes and the blue ones represented for down-regulated genes. (B) The upset plot showing the number of all of unique and some of shared genes of hippocampal cell types. (C-F) The biocprocess in GO terms enriched separately by up or down-regulated genes of astrocyte, endothelia, microglia and oligodendrocyte. Criteria for DEGs by Muscat: p_value < 0.05, abs(log2FC) > 0.25).

Similarly, endothelial cells showed 565 DEGs (322 upregulated and 243 downregulated) (Fig.3.A), of which 268 were cell type-specific. Notably, a subset of 116 DEGs was shared between these two cell types (Fig.3.B).

These DEGs were subsequently categorized based on their direction of regulation (upregulated or downregulated) and subjected individually to pathway enrichment analysis using Gene Ontology (GO), in order to elucidate their functional roles and the molecular basis of METH-induced cognitive decline. The upregulated and downregulated Biological Processes (BP) from the GO analysis are presented (Fig.3.C – F and SI.2). Moreover, analysis of cell type-specific enrichment patterns revealed that, in addition to distinct functional effects, there exist complex co-regulatory mechanisms across different cell types in response to chronic METH exposure.

Neuroinflammation is a complex process that can be triggered by various factors, such as exposure to toxic substances like METH. METH-induced neuronopathies, in particular, have been found to constantly produce a range of impact factors and cells’ changes that contribute to the exacerbation of neuroinflammation. Microglia up regulates immunoprotein receptor genes (such as *Fcer1g*, *Fcgr3* and *Il1rl2*) to enhance immune sigal reception and cell chemotaxis (*Ccl2*, *Cxcl14*, *Trpv4*, et al.) (Fig.3.E). Astrocytes enhance their capacity for gliogenesis and migration through the expression of key regulatory genes (*P2ry12*, *Rras*, *Efemp1*,et al.) thereby modulating the inflammatory response via the NF-κ B signaling pathway (involves *Tnfaip6*, *Nfkb1*,*Nfkbia*) (Fig.3.C). In addition to the responses of neurogliocytes, genes like *Grn* and *Trem2* (associated with astrocye and microglia activation) are up-regrated in both microglia and T cell, suggesting potential communication between central resident and peripheral immune cells, which may jointly participate in the neuroinflammatory response. T and NK-T cells become more active in leukocyte or lymphocyte mediated immunity, but T cells active by complement-mediation (*C1qb*, *C1qc*, *C1qa*, *Trem2*) while NK-T cells depend more on reception of cytokines (*Il18r1*, *Fcer1g*) or recognition of receptors on membrane (*Itgb2*, *Ptprc*, *Tyrobp*,et al.). However neutrophils down-regulate chemokine-mediated signaling pathway (*Cxcr4*, *Ccl3*, *Ccl4*) that may indicates neuroinflammation changes from acute (leukocyte-leading) to chronic (lymphocyte-leading). Macrophage also weaken leukocyte activation (*Cd28*, *Itgam*, *Il18*, *Tnfsf9*, et al.) but strengthen T cell activation.

The maintenance of a stable neural microenvironment relies on a healthy blood-brain barrier (BBB) system(13). METH can induce structure and function disruptions in the BBB, although its chronic effects remain uncertain. In our study, we observe that endothelial cells upregulate detoxification funtions (*Gstm1*, *Abcg2*, *Fbln5*, et al.) and autophagy-related genes (*Qsox1*, *Fnbp1l*, *Trp53inp1*, et al.), while downregulating angiogenesis-associated genes (*Hipk2*, *Tnfrsf1a*, *Vegfc*, *Cldn5*, et al.) and cell-cell adhesion molecules (*Hmcn1*, *Ceacam1*, *Cdh10*, et al.). Furthermore,endothelial dysfunction is also implicated in learning or memory deficits and cognitive impairments (*Dbi*, *Bche*, *Psen2*, *Prnp*, et al.) as well as nervous system development (*Macf1*, *Ptprz1*, *Vegfc*, *Ctnnb1*, et al.) suggesting an interplay between neuronal activities and vascular functions (Fig.3.D). Similarly,dysfunctions in *Cldn5^+^* mural cells are associated with altered vascular permeability (*Ctnnbip1*, *Pde3a*, *Ceacam1*, *Cldn5*) and developmental processes (*Atp2b4*, *Sfrp2*, *Vegfc*, *Tgfbr2*, et al.), as well as cognitive impairments (*Psen2*, *Bche*, *Nf1*, *Prkn*, et al.), and increased autophagic cell death (*Bmf*, *Bnip3*, *Lamp1*, *Cdkn2d*) after METH trearment. As a structure of arteriole in hippocampus, SMCs exhibit suppressed inflammatory responses such as leukocyte migration (*Abl2*, *Fcer1g*, *Vcam1*, *Icam1*, et al.), glial cell activation (*Syt11*, *C1qa*, *Cx3cr1*, *Trem2*, et al.) and IL-6 production (*Dhx9*, *Fcer1g*, *Syt11*, *Il6ra*).

The response of BBB cell types to METH is heterogeneous(14). In our data, within the Wnt signaling pathway—which promotes angiogenesis and vascular remodeling(15)—both *Cldn5*^+^ and *Cldn5*^−^ mural cells show downregulation through distinct sets of DEGs, whereas endothelial cells exhibit upregulation of this pathway. Cell-substrate adhesion is enhanced in mural cells and SMCs, but reduced in ECs. Although this ultimately results in increased BBB permeability, the underlying mechanisms differ between the inner and outer layers of the vasculature.

Metabolite shuttling and extracellular vesicle signaling provided by oligodendrocytes are essential for the normal functioning of neurons, particularly their axons(16). In our study, we observed significant changes in the metabolic status of oligodendrocytes (Fig.3.F), including alterations in organic acid biosynthesis (*Cyp27a1*, *Per2*, *Glul*, et al.), amino acid metabolic (*Glul*, *Bcat1*, *Hmgcll1*, et al.), glycolipid metabolic process (*Pikfyve*, *Pigc*, *Pnpla7*, et al.) and other processes. Additionally, METH influences neural structure and function-related changes such as axonogenesis (GO terms:axonogenesis, postsynapse organization, central nervous system neuron development, et al.) and synaptic vesicle signaling (GO terms: vesicle-mediated transport in synapse, synaptic vesicle recycling, vesicle localization, et al.), which are partially upregulated or downregulated. Furthermore, oligodendrocyte precursor cells exhibit negative effects on cell development and synaptic plasticity while also increasing neuroinflammation response to DNA damage stimulus. These effects may contribute to neuronal dysfunction and ultimately lead to cognitive decline. Astrocytes play a vital role in supporting neuronal function throughout the central nervous system by regulating synapse development and neurotransmitter release at synapses while providing adequate energy supply and modulating information processing within neurons(17). During chronic METH abuse, astrocytes downregulate bioprocesses involved in translation and metabolism (Fig.3.C), such as translation at synapse (*Rpl12*, *Rpl35*, *Rps26*, et al.) and the generation of precursor metabolites and energy (*Adh5*, *Ndufa5*, *Bpgm*, et al.), which may impede normal neurotransmitter vesicle transmission. Simultaneously, astrocytes enhance bioprocesses related to multiple inorganic cation transportation including potassium (*P2ry12*, *Kcnd3*, *Slc24a2*, et al.), sodium (*Slc24a2*, *Nkain1*, *Hcn2*, et al.), or calcium (*P2ry12*, *Tmem38a*, *Slc24a4*), which may interfere regulation of neural membrane potential. However, we also observe that astrocytes have positive effects of neuron projection extension (*Nrp2*, *Rtn4*, *Jade2*, et al.), axon ensheathment (*Abca2*, *Mal*, *Ugt8a*, et al.), or oligodendrocyte differentiation (*Abca2*, *Mal*, *Mag*, et al.). These results may imply that astroctyes have dfferent subsets performing diverse functions according to their positions and cellular contacts in hippocampus.

Adult hippocampal neurogenesis (AHN) is a complex and dynamic developmental process regulated by a multitude of factors. It involves a sequential progression from neural stem cells (NSCs), through radial glia-like cells and neuroblasts, to fully differentiated mature neurons. This process plays a crucial role in cognitive functions such as learning, memory formation, and emotional regulation. Furthermore, alterations in AHN have been associated with memory impairments, highlighting its significance in maintaining hippocampal plasticity and overall brain function(18). At the initiation of adult hippocampal neurogenesis (AHN), we observe that neural stem cells (NSCs) downregulate pathways related to the generation of precursor metabolites and energy (e.g., *Sdhaf4*, *Nduf* family, *Uqcrh*, *Gapdh*), suggesting that METH may impair the proliferation and differentiation potential of NSCs. Furthermore, METH induces endoplasmic reticulum stress (e.g., *Tmco1*, *Atf3*, *Hspa5*, *Gsk3b*), activates apoptotic processes (e.g., *Btg2*, *Prnp*, *Cebpb*, *Ctsz*), and promotes autophagy (e.g., *Ddrgk1*, *Ctsa*, *Mcl1*, *Sh3glb1*) in NSCs. These alterations consequently lead to increased expression of β-amyloid-related genes (e.g., *Prnp*, *Apoe*, *Psenen*, *Rtn4*) and IL-6-related genes (e.g., *Ncl*, *Fcer1g*, *Il1a*, *Lbp*), which may trigger neuroinflammatory responses. Adult NSC activation in mice is known to be regulated by circadian rhythms and intracellular calcium dynamics(19). In our dataset, astrocytes from METH-treated mice show enhanced expression of circadian rhythm-related genes (e.g., *Per2*, *Ptgds*, *Ciart*, *Id3*), calcium-mediated signaling pathways (as previously mentioned), and genes involved in the regulation of neurogenesis (e.g., *Per2*, *Mag*, *Fbxo31*, *Slc7a5*). These findings suggest that astrocytes may attempt to compensate for METH-induced damage by activating NSCs; however, this compensatory mechanism could accelerate the depletion of the NSC pool.

During the neuroblast stage, METH exposure increases oxidative stress through upregulation of genes associated with oxidative stress response (e.g., *Setx*, *Prkra*, *Agap3*, *Lig1*) and alters DNA regulatory mechanisms involved in cell division (e.g., *Ciz1*, *Rpa2*, *Mcm7*, *Ssbp1*). Additionally, METH suppresses cell-substrate adhesion by downregulating genes such as *Myoc*, *Ccn1*, *Ptn*, and *Lrp1*. As the remaining neuroblasts mature into neurons, METH exerts more pronounced effects, resulting in a higher number of DEGs. Under METH exposure, neurons derived from NSCs upregulate pathways related to endoplasmic reticulum stress (e.g., *Tmco1*, *Atf3*, *Hspa5*, *Pdia3*), TNF superfamily cytokine production (e.g., *Ncl*, *Fcer1g*, *Il1a*, *Lbp*), neuron death (e.g., *Prnp*, *Ctsz*, *Mcl1*, *Jun*), leukocyte chemotaxis (e.g., *Fcer1g*, *Lbp*, *Cxcl10*, *Hmgb1*), and β-amyloid metabolic processes (e.g., *Prnp*, *Apoe*, *Psenen*, *Rtn4*). Simultaneously, these neurons downregulate key pathways involved in energy metabolism (e.g., *Sdhaf4*, *Nduf* family, *Cycs*, *Gapdh*) and mitochondrial structure and function (e.g., *Slc25a* family, *Vdac1*, *Atpif1*, *Acaa2*).

Cajal-Retzius cells (CRs) are transient neurons that almost completely disappear in the neocortex after birth but survive up to adulthood in the hippocampus at a certain percentage. CRs are synaptically integrated into hippocampal circuits relating to hippocampal-related behaviors and network. Abnormal survival of CRs has been linked to deficits in adult neuronal and cognitive functions(20, 21). In our analysis, CRs down-regulate various neural growth and maturation-relevant processes including axon extension (e.g.,*Sema6d*, *Dclk1*, *Cttn*, *Sin3a*), dendrite genesis (e.g., *Nlgn1*, *Dclk1*, *Farp1*, *Il1rapl1*), synapse organization (e.g., Lrrtm4, Lhfpl4, Farp1, Mark2) and neuron migration (e.g., *Nr4a2*, *Dclk1*, *Sema6a*, *Mark2*, *Cdkl5*).

In summary, METH exerts direct neurotoxic effects on NSCs and their developmental trajectory, while also indirectly disrupting the function of neurogenesis-supporting cells, ultimately contributing to cognitive decline.

Ependymal cells, a type of neuroglia that provides support to neurons, form the epithelial lining of the brain ventricles(22). METH exposure induces significant changes in ependymal cells, including the upregulation of genes associated with cilium organization and motility (e.g.,*Meig1*, *Cfap* family, *Lca5*, *Tmem216*), suggesting enhanced cerebrospinal fluid circulation and a potential compensatory response to toxic substances by facilitating the clearance of harmful molecules. However, METH also downregulates genes involved in intracellular transport (e.g., *Ptpn14*, *Pdcd10*, *Txn1*, *Sorl1*) and protein processing (e.g., *Gsn*, *Txn1*, *Tmsb10*, *Sorl1*), indicating impaired cellular functions that may compromise ependymal cell homeostasis.

Fibroblasts, which originate from embryonic mesenchymal cells, perform a variety of complex and diverse functions in the hippocampus, including extracellular matrix (ECM) homeostasis, secretome regulation, mechanosensation, progenitor cell activity, and immune modulation(23). However, their specific roles in this brain region remain incompletely understood. Compared to the saline group, METH-treated fibroblasts exhibit increased expression of genes associated with responses to metal ions (e.g., *Gsn*, *Mt2*, *Mt1*, *Id2*, *Hnrnpa1*) and toxic substances (e.g., *Gstm1*, *Inmt*, *Mt2*, *Mt1*), as well as markers linked to astrocyte differentiation (e.g., *Vim*, *Plpp3*, *Sox9*, *Id2*). Notably, downregulation of the JNK and MAPK signaling pathways (e.g., *Map4k4*, *Sh3rf1*, *Ccn2*, *Gadd45b*, *Dab2*) is associated with reduced fibrosis, suggesting that fibroblasts may undergo functional transitions— from stress responses such as cell death and inflammation—to roles in cellular support and transformation.

### 4. METH-induced alterations of crosstalk between cell types in hippocampus led to hippocampal environment disturbance

Considering the neurotoxic effects of METH on various cell types in the hippocampus and its intricate impact, we conducted an investigation into hippocampal intercellular communication to elucidate the role of these interactions in cognitive decline. Our findings reveal a slight increase in both the total number of cell communication interactions (from 18,396 to 19,054) and their strength (from 0.883 to 0.841) following METH treatment (Fig.4.A). The results demonstrate that chronic METH abuse induces a greater diversity of changes in the differential number of interactions among cell types compared to changes in interaction strength (Fig.4.B and C). For instance, astrocytes mildly enhance the number of interactions with mural cells, ECs, and OPCs but decrease the strength of these interactions, suggesting reduced efficacy of communication signals under METH influence. Neuron derived from NSCs increase the number of interactions with most cell types while exhibiting minimal changes in interaction strength, which corresponds to the up-regulation of TNF superfamily cytokines and damage signals associated with ER stress or amyloid-beta mentioned in DEGs analysis. In association with neuroinflammatory response, microglia significantly augment their own interaction strength, potentially contributing to neuroinflammation.

**Fig. 4.**
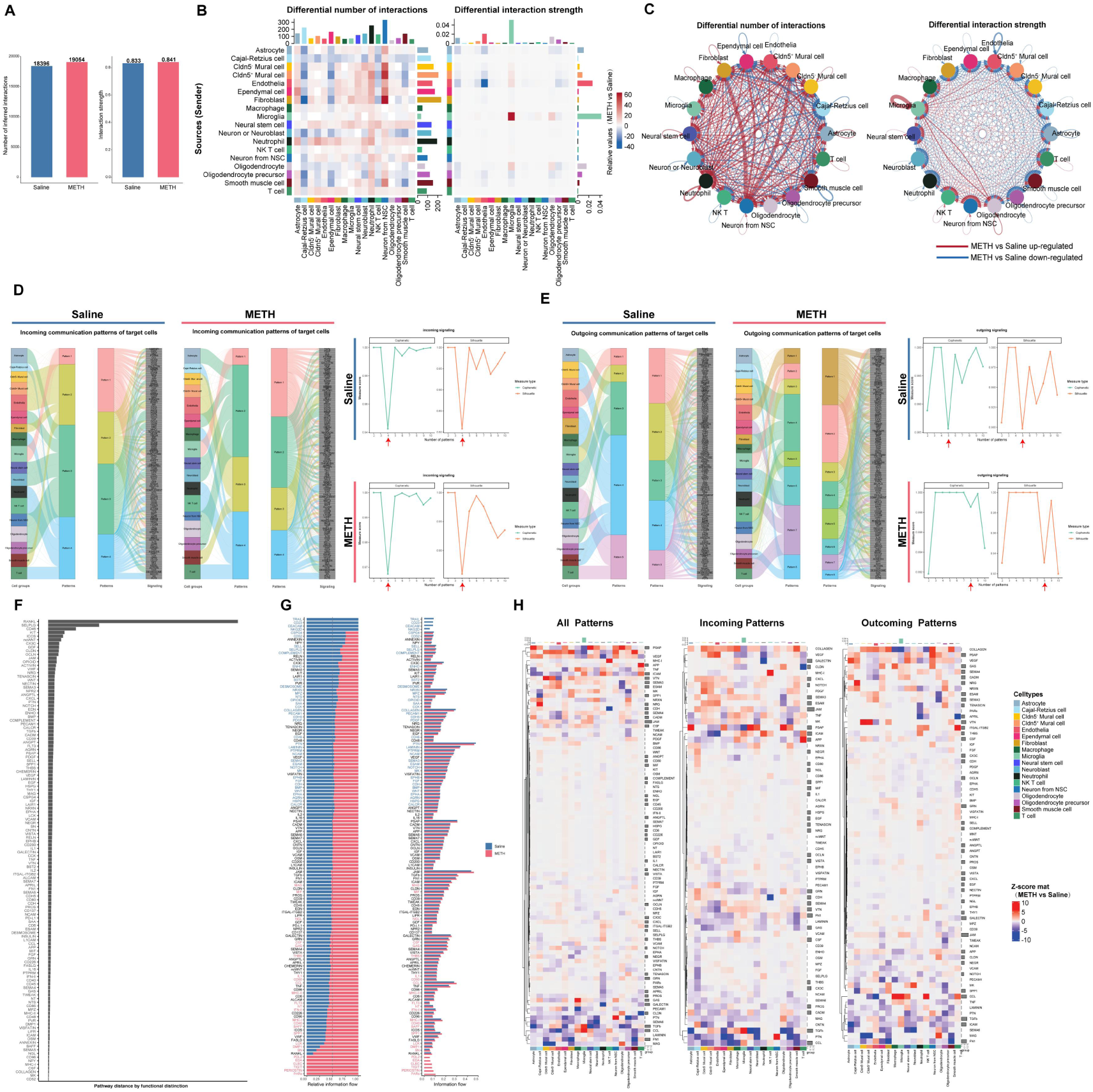
Hippocampal cell types crosstalk alterations induced by chronic METH treatment. (A) The total number of cell communication interactions and the intensity of interactions in saline and METH groups. (B-C) The comparison of the number and the strength of interactions of each cell types between METH and saline groups. (D-E) The incoming and outgoing communication patterns of secreting cells of each groups and the inflection point parameter in clustering algorithm. (F) The identificaton of signaling networks with large (or small) differences according to the Euclidean distance in a shared two-dimensional space. (G) The comparison of the overall information flow of each signal path between two groups. (H) Heatmap showing the incoming, outgoing and all communicated informations’ comparison of each signal of each cell type.

From the perspective of cellular communication patterns (incoming and outgoing), hippocampal cells treated with METH exhibit 4 incoming patterns similar to the Saline groups, primarily related to microglial activation, BBB function, neural development, and immunity (Fig.4.D). However, in terms of outgoing patterns, the communication patterns of METH groups are divided into 8 distinct patterns compared to 5 in the Saline group. These outcoming patterns mainly involve microglia activation, endothelial cell signaling, neural development processes, immune responses, as well as a mixed pattern reflecting changes in the neural microenvironment (Fig.4.E). The disruption caused by METH results in confusion within cellular crosstalk specifically through disturbances in outcoming communication pathways within the hippocampus. To further elucidate how METH affects cellular communication networks concretely, we conducted an analysis focusing on specific signal pathway alterations. By calculating Euclidean distance (Fig.4.F), we identified several key differential functions between METH and Saline groups: RANKL (Receptor Activator of Nuclear Factor-κ B Ligand, belongs to tumor necrosis factor (ligand) superfamily, also proteinically known as TNF-related activation-induced cytokine), SELPLG (proteinically known as p-selectin, involving in leukocyte and microglia activation(24)), and CD46 (proteinically known as complement regulatory protein, as a membrane cofactor mediated cleavage cofactor of C3b and C4b, which is a key regulator of classical and selective complement activation cascade, as well as inflammation(25)). We then compared overall information flow across each signal path to determine conservative and METH-specific signaling pathways (Fig.4.G). Additionally, we also examined changes occurring within each outgoing or incoming signal pathway for individual cell types (Fig.4.H).

### 5. Pseudo-time series analysis revealed that METH caused changes in hippocampal neurogenesis in mice

Adult neurogenesis in the dentate gyrus of the hippocampus plays a crucial role in cognitive function in mice and is influenced by the local microenvironment or niche(26). However, limited evidence exists regarding the impact of METH exposure on the capacity of NSCs to differentiate into neurons, astrocytes, oligodendrocytes, and other cell types. To address this, we conducted pseudotime analysis to explore how chronic METH exposure affects NSC differentiation trajectories.

Along the temporal axis, with NSCs as the starting point, we observed that under normal conditions (Saline group), NSCs (cluster 20) predominantly differentiated into neuroblasts (cluster 17) compared to astrocytes (clusters 6 and 27), following a typical neural developmental trajectory (Fig.5.A). In contrast, METH exposure disrupted this pattern, showing an increased tendency for NSCs to differentiate into astrocytes rather than neuroblasts (Fig.5.D), suggesting impaired neurogenesis.

**Fig. 5.**
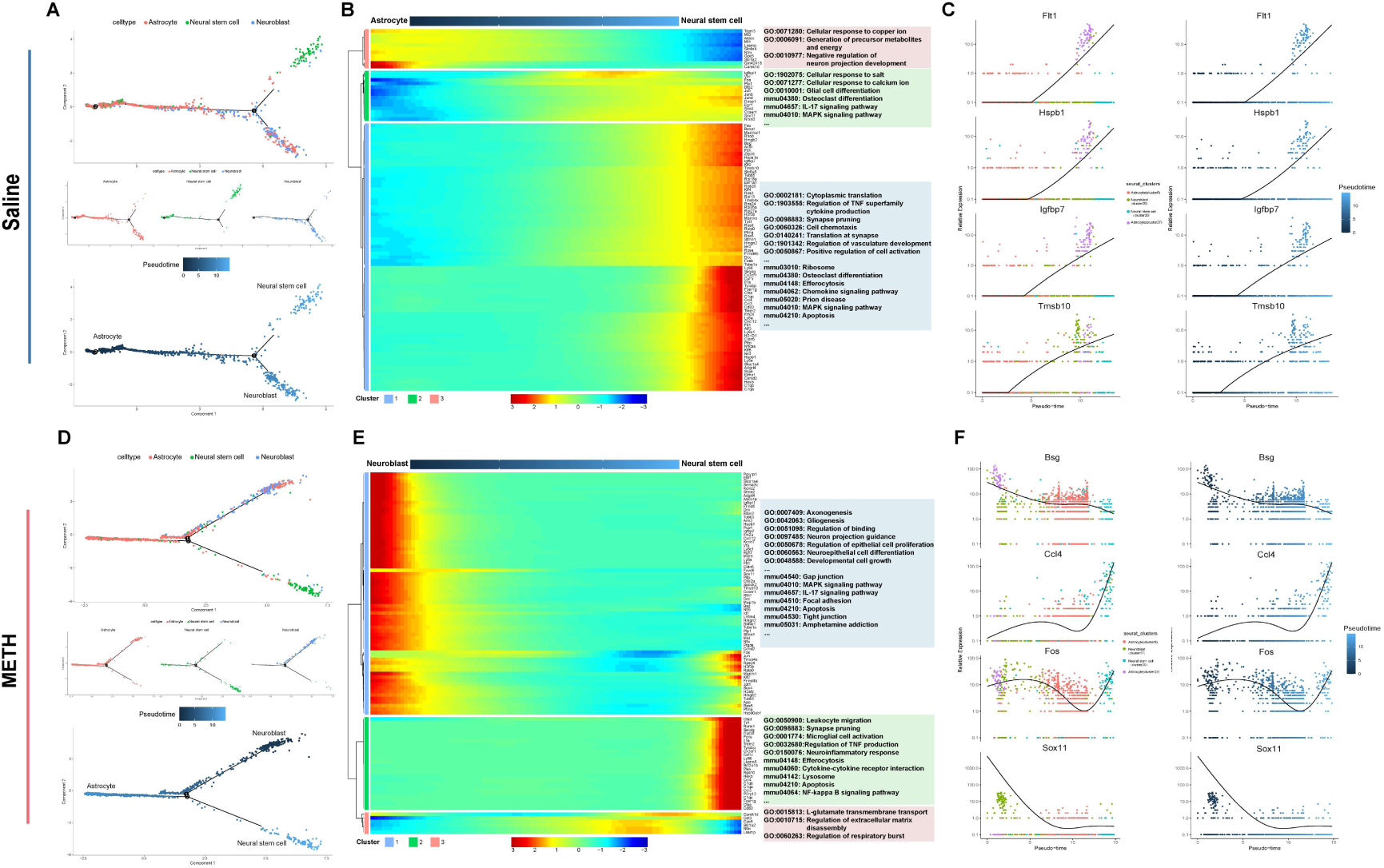
The development direction of hippocampal neural stem cells and the regeneration of hippocampal nerve were analyzed in pseudo-time series. (A, C) The pseudo temporal differentiation locus of neural stem cells, astrocytes and neuroblasts in saline (A) or METH (C) group. (B, D) Heatmap showing the expression of specific genes(rows) in subclusters (columns) along the maturation trajectory from neural stem cells to neuroblasts or astrocytes in saline (B) or METH (D) group, and GO or KEGG funtional enrichment of those genes. (E, F) Pseudotime expression graphs of 4 representative specific marker genes in these cell types showing the development tendency difference in saline (E) or METH (F) group.

Our analysis reveals that the altered neural developmental trajectory in the METH group is associated with changes in the expression of genes such as Bsg, Ccl4, Fos, and Sox11 (Fig.5.F), as illustrated by other differentially expressed genes in the heatmap (Fig.5.E). In contrast, normal developmental progression from NSCs to neuroblasts is more closely associated with the expression of Flt1, Hspb1, Igfbp7, Tmsb10, and other related genes (Fig.5.B and C).

### 6. The change of transcription factor regulatory network is also the cause of cognitive decline caused by METH

We conducted an analysis of the differential expression of transcription factors in each cell type between METH and Saline mice (Fig.6.A and B). Subsequently, we constructed networks of transcription factor-target genes to predict changes in cellular functions (Fig.6.D-H, SI.3).

**Fig. 6.**
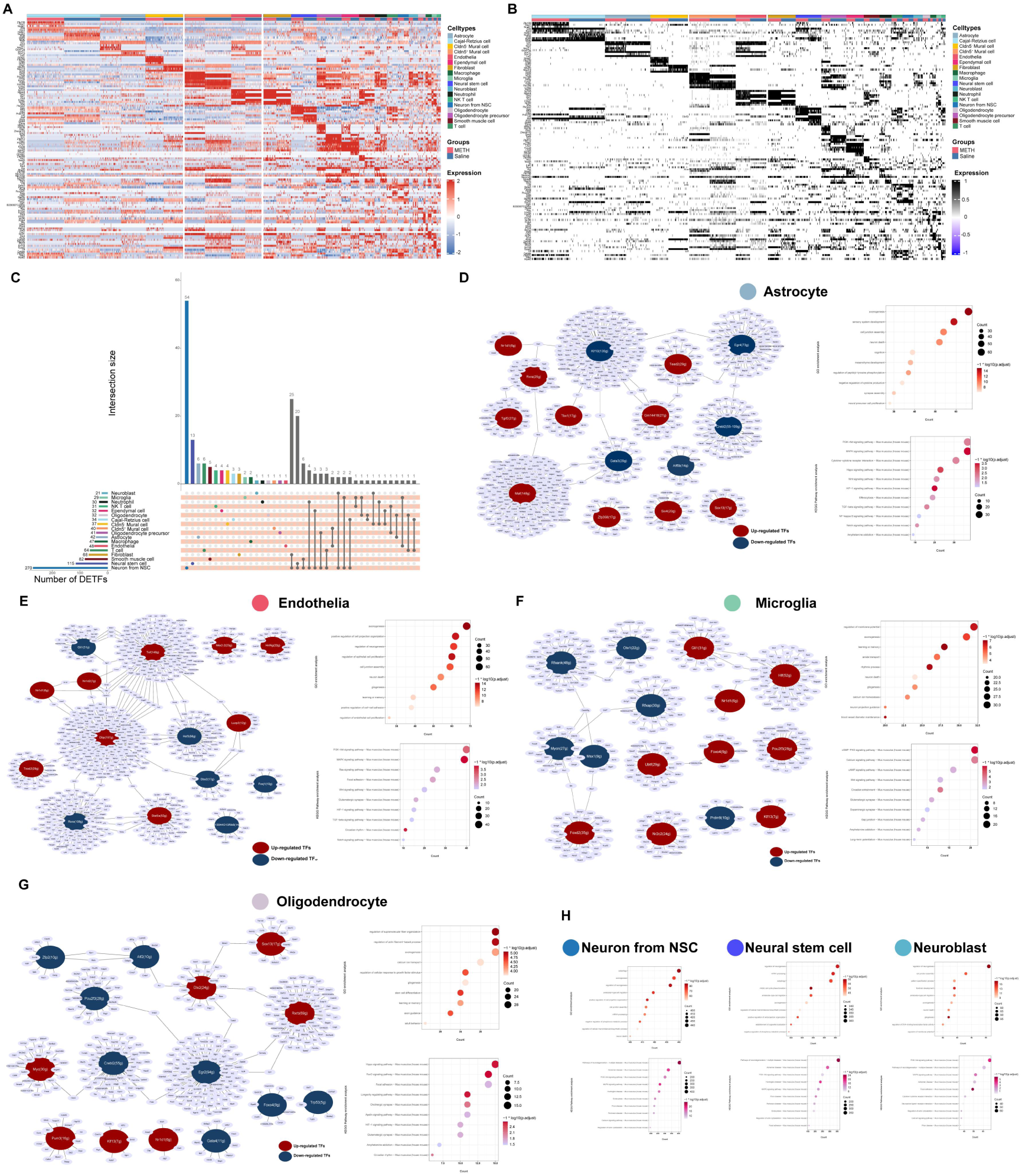
Transcription factor (TF) analysis and function prediction. (A-B) Heatmap of the average expression and the area under the curve scores of TF motifs estimated per cell by SCENIC. Shown are top five differentially activated motifs in each cell type of saline and METH groups, respectively. (C) The upset plot showing the number of all of unique and some of shared TFs of hippocampal cell types. (D-G) Gene regulatory network analysis and function enrichment using top 10 of SCENIC identifies critical TFs of astrocytes, endothelia, microglia and oligodendrocytes. (H) Enrichment analysis of target genes predicted by TFs of neuronal development related cell types (Neuron from NSC, NSC and neuroblast).

In astrocytes, a total of 42 differentially expressed transcription factors (DETFs) were identified (Fig.6.C), and their associated target genes were enriched in biological processes related to neurogenesis and development, cognitive function, regulation of inflammatory responses, as well as signaling pathways including PI3K-Akt, MAPK, and cytokine-cytokine receptor interactions, among others (Fig.6.D).

Similarly, endothelial cells exhibited 48 DETFs, whose target genes were also enriched in biological processes related to neural development, including axonogenesis, neuronal differentiation, and functions associated with learning and memory. Endothelial cells share common signaling pathways with astrocytes, such as PI3K-Akt and MAPK (Fig.6.E). Furthermore, biological processes specifically associated with endothelial cells, such as endothelial cell proliferation, cell-cell adhesion, and cell junction assembly, were found to be involved in signaling pathways including Ras, focal adhesion, and Wnt, among others.

Although only 29 DETFs were identified in microglia, they play a significant regulatory role in processes such as membrane potential modulation, axonogenesis, and neuronal death. These biological processes are closely associated with synaptic pruning and neuronal survival, which may ultimately contribute to cognitive dysfunction. Signaling pathways including calcium, Wnt, cGMP, and cAMP are actively involved in the aforementioned processes (Fig.6.F).

As key supporters of neuronal bodies and axons, oligodendrocytes exhibit 32 DETFs that regulate processes related to cytoskeletal fiber organization and axon development. This suggests that METH exposure disrupts the transcriptional regulation of genes involved in neural support, thereby increasing neuronal vulnerability to damage. These effects may be mediated through signaling pathways such as Hippo, FoxO, Apelin, and Focal Adhesion. Whether the transcriptional disruption affects neuronal growth itself or the maintenance of surrounding cells and the neural environment under chronic METH treatment remains an important factor contributing to cognitive decline in mice.

Interestingly, neurons derived from NSCs exhibit the highest and second-highest numbers of DETFs, respectively, and share highly similar functional enrichment terms among their target genes. These terms are primarily associated with neurodegenerative processes such as autophagy and neuronal death, axonogenesis, ameboid cell migration, and biosynthetic activities, which are also comparable to those observed in neuroblasts (Fig.6.H). This indicates that METH may exert common neurotoxic mechanisms in both mature neurons and neural precursor cells by modulating similar pathways through the expression of differentially regulated neural transcription factors.

## Discussion

As a psychostimulant drug widely use in worldwide, METH can elevate the risk to various neurological neuropsychiatric disorders due to its addictive and neurotoxic effect. In our study, animal behavior experiments illustrate that chronic METH abuse can cause cognitive decline in mice, which is manifested in spatial cognition, working memory and new object learning. Although previous studies also have similar concusions of METH-induced learning and memory impairments and put forward a variety of therapeutic improvement schemes (e.g. melatonin, oxytocin, housing and tetrahydropalmatine)(27–30), it seems that there are few practical, effective, and recognized treatments for chronic METH abuse-induced cognitive dysfunction therapy. Because researchers have found that METH can affect many organs throughout the body including most areas of the brain by complex and diverse molecular mechanisms, mainly involving excitatory toxicity, neurotransmitter disruption, oxidative stress, cytotoxicity (apoptosis, necrosis, and autophagy), inflammatory response, metabolism disorder et al(31–33). Furthermore, unlike acute METH-induced injury, chronic METH abuse represents a more complex process characterized by concurrent neurotoxic damage and reparative responses(34–36).

The hippocampus plays an essential role in a range of advanced brain functions, including memory and emotional regulation, and spatial navigation, and its impairment or dysfunction have been proved to closely associate with neuropsychological diseases like neurodegeneration, affective disturbance, and learning disabilities(37).

On the base of our animal behavior results, we used scRNA-seq to investigate all mouse hippocampal cell types except for mature neurons and to explain the heterogeneity of METH effects between different types of cells and provide more targeted mechanisms for METH-induced neurotoxicity and targets for precise therapy. Given that this study aims to capture the complete mRNA expression profile of intact cells, we employed scRNA-seq technology. In contrast to single-cell nucleus sequencing (snRNA-seq), this approach necessitates that cells be maintained under conditions as close to physiological states as possible, thereby requiring high structural integrity, robust viability, and minimal exposure to external perturbations. Accordingly, during sample collection and the preparation of single-cell suspensions, we processed the tissues as rapidly and gently as feasible. Nevertheless, mature neurons — due to their highly differentiated state, limited deformability, and the presence of long axons — are particularly susceptible to damage or functional inactivation during the dissociation process, and are therefore frequently excluded prior to sequencing. Therefore, we propose that the neurons captured in this study predominantly represent immature neurons or a distinct subpopulation of neurons with relatively smaller cell bodies. Although we did not detect the expression profiles of mature neuronal genes, our dataset provides a comprehensive and detailed characterization of various non-neuronal cell types. Each analyzed cell contains the complete mRNA expression profile, which accurately reflects its functional state. Accumulating evidence indicates that non-neuronal cell populations in the brain play a critical role in chronic neurodegenerative diseases, with their functional states significantly influencing neuronal activity and serving as potential early biomarkers, contributors to disease pathogenesis, and therapeutic targets(38, 39). Notably, the cognitive impairments induced by chronic METH exposure exhibit certain pathological similarities to those observed in Alzheimer’s disease and Parkinson’s disease(40, 41). Our dataset offers valuable insights into the functional alterations, predictive dynamics, and intercellular communication of non-neuronal cells following chronic METH exposure, thereby providing robust biological information to enhance our understanding of METH-induced neurotoxicity and cognitive dysfunction.

### 1. Revealing a novel pattern of neuronal death and elucidating the impact of neuronal interactions with other cells during development and maturation

METH can cause abnormal neuronal function in various direct or indirect ways. METH has a direct toxic effect on neurons causing neuronal damage by various programmed cell death, such as apoptosis, autophagy, necroptosis, pyroptosis, and ferroptosis(42). In spite of mature neurons loss during dissociation, we still isolated the neurons remaining a few NSC markers in some degrees and we considered these neurons developed from NSCs on account of that adult hippocampal neurogenesis is a continuous process. Consistent with previous results, we found a few genes of these neurons related to multiple neuron death biological processes exist differences. For example, Cebpb mediate endoplasmic reticulum stress by Nupr1-C/EBP homologous protein (CHOP) pathway, Prnp/Psenen/apoe are associated with neurodegeneration disease in amyloid-beta metabolic process, Gsk3b increases phosphorylation of Tau, Ddrgk1/Mcl1/Sh3glb1 play important role in mitochondrial autophagy, ATF3 forms dimer with c-Jun to promote apoptosis(43). We also noticed TNF superfamily cytokines and ILs overexpression in these neurons may attracts microglia or actives astrocytes leading to neuron death. Abnormal neuronal function is also indirectly related to METH’s effects on various auxiliary cells. Our cellchat data showed these neurons generating more communication signals to vascular cells, fibroblasts and ependymal cells after chronice METH treatment. As sources, these neurons upregulate signals like Vegf (Vascular endothelial growth factor, promoting blood vessel growth and increasing permeability.), Sema3b and Sema3c(are critical for neuron projection, guidance of axons, dendritic spine pruning and ECs repulsion(44–46)), Ptn and Mdk (Pleiotrophin and Midkine, as neurotrophic factors, is critical in different steps of differentiation of different cells both in development and in wound repair, especially in neural regeneration, and is up-regulated in drug abuser and neurodegenerative diseases patient(47).), Igf2 (Insulin-like growth factor 2, emerged as a critical molecule of synaptic plasticity and learning and memory(48).), and Cxcl12 (concerning about regulating generation, positioning and maturation of new neurons(49, 50), as well as the communication with blood vessels(51).), but downregulate signals like Bmp6 (Bmp6 level shows an increase in AD-related neurogenic deficits(52), and critical functions to EC differentiation(53).), Bmp7 (Bmp7 expression in radial glial cells may promotes neurogenesis, inhibits gliogenesis(54).), and Fgf2 (a crucial molecule modulating cell proliferation and survival in central nervous system(55).). As receptors, there neurons recept signals from vascular cells mainly about extracellular matrix, such as Col (collagen) families and Lam (laminin) families. Our analysis showed METH impedes the neuronal maturation process which closely relied on the regulation of vascular cells (Fig.7).

**Fig. 7.**
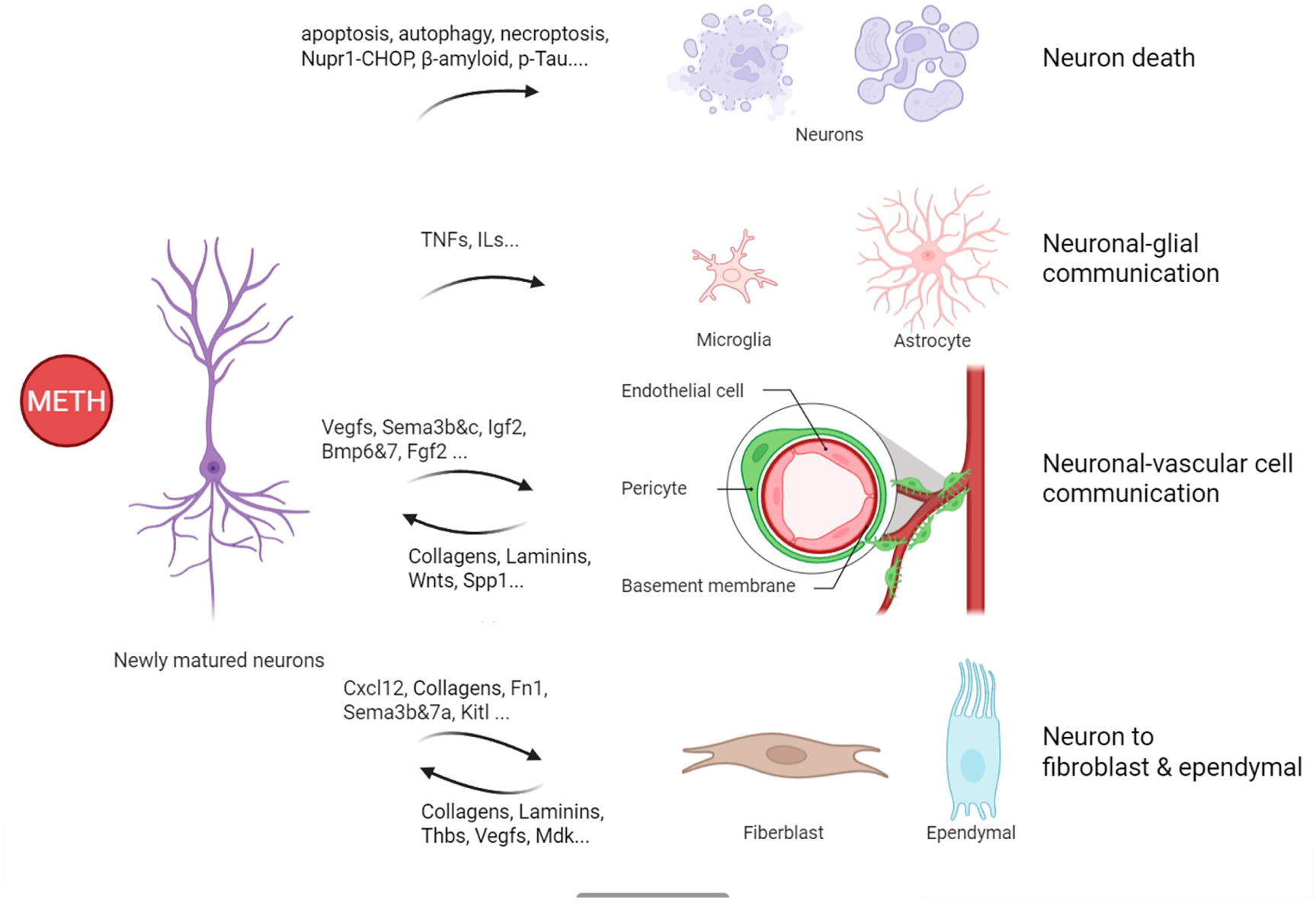
A schematic illustration of the effects of METH on neuronal cell death during development and maturation and its interactions with glias, vascular cells, fiberblast and ependymal.

### 2. Analysis of dynamic changes in neurogenesis to track spatiotemporal shifts in cell composition, signaling, and functional integration during nervous system development and regeneration

METH can cause damage to neurons through various mechanisms, while also have impacts on neural stem/progenitor cells. However, these impacts seem to be different between researches of actue or chronic abuse, intermittent or continuous administration, addiction or withdrawal(56–59). In the researche of chronic METH abuse, it has reported that METH mainly has negetive effects on stem/progenitor cells resulting decreasement proliferative and neurogenesis capacity(56, 60), and even cell death(61, 62). Here our analysis provided in vivo evidences of neurogenesis alterations under chronice METH abuse. We confirmed METH’s neurotoxicity to NSCs and progenitor cells like neuroblasts and CRs, and ulteriorly we used pseudotime to explore its influences on differentiation process and dynamics of neural stem cells. The variations of characteristic genes on each branches of neural development trajectory may reveal inflection points in the direction alterations of NSC development under METH effect. It is also necessary to emphsize accurate quantifications of various types of cells counting in the neurogenetic locus will be helpful to lucubrate the toxicity of METH on neurogenesis (Fig.8).

**Fig. 8.**
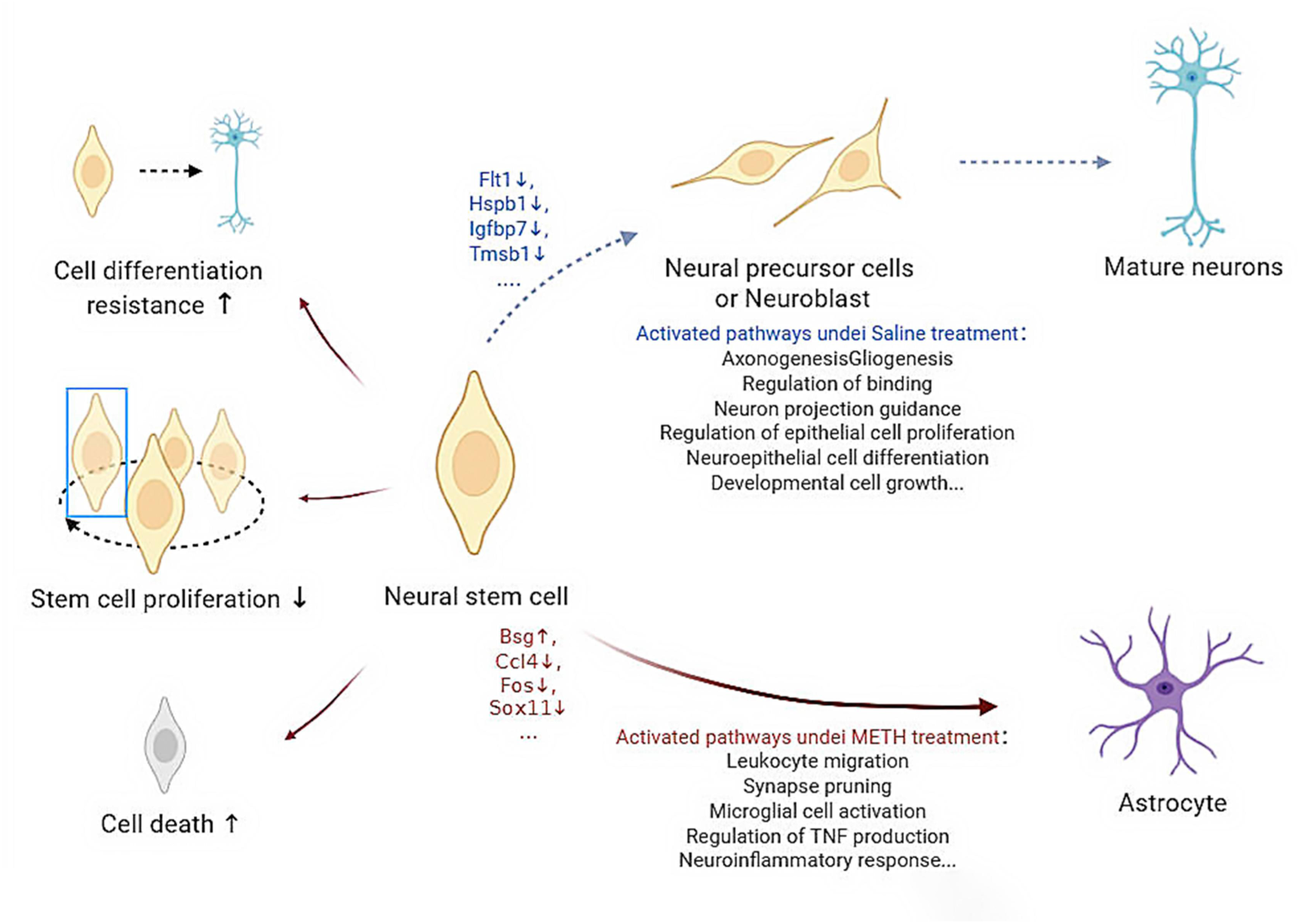
A schematic illustration of the effects of METH on cell proliferation capacity and cellular activity in neural stem cells, as well as associated alterations in gene expression and signaling pathways during their differentiation into neurons or astrocytes.

### 3. Establish a comprehensive cellular interaction map of the neurovascular unit

BBB injury is one of neurotoxic mechanisms of METH neurotoxicity, however, the impact of METH on BBB and its consequent effects on neural function far exceed the barrier dysfunction and abnormal substance transport mentioned in current researches. Previous studies of METH neurovascular toxicity have reported that ECs structure and function disorder and astrocytes-released cytokines or damage factors induced injury are parts of proper mechanisms for BBB injury(14), however these studies usually consider a single aspect of cell type or mechanism within BBB while current researches emphasize that neurovascular unit (NVU) comprises more informations about microcirculation integrity, vascular function and cellular cooperation, which are important to cognition regulation(63). In our study, we parsed most of NVU components which are consisted of ECs, mural cells, SMCs, gial cells (astrocytes, oligodendrocytes and microglia), neurons and basal matrix. We found that ECs can be grouped into several subsets may related to the difference between arteries and veins and the grade of vascular branches, which make it possible to anaylze endothelial dysfunctions like insufficient nutrient and oxygen transport, low clearance efficiency of metabolic waste in cerebral circulation, or leukocyte adhesion and migration. Moreover, mounting evidences correlate mechanical stress with endothelial function, such as barrier function, inflammatory signaling, apoptosis, oxidative stress, endothelial mesenchymal transformation and aging(64), which means ECs located in different fluid states of vascular segments may be regulated variously as results of METH-induced cerebral blood flow abnormalities(65–67). In the meanwhile, the influence of the endothelium on other cell types is extensive, including but not limited to vascular tone or structure, maintenance of collateral vessels, neurotransmission and neurogenesis(63), as our DEGs analysis found that the effects of METH on ECs are not only to damage the junction structure and cause autophagy apoptosis, but also to change the association with other cells, such as GO terms: vesicle-mediated transport in synapse, neurotransmitter transport, or glial cell proliferation. Although there are few METH-induced neurovascular toxicity researches concerning mural cells, who are thought to include pericytes, SMCs and fibroblasts(68), we tried to distinguish them by expression levels of maker genes and eliminate the effect of doublets, and we interestingly found that they share many common genic features as ECs and SMCs, of which some of mural cells have similar expression levels of marker genes as ECs especially. As a result, we classified mural cells as *Cldn5*^+^ and *Cldn5*^-^ mural cells by the similarity to ECs, and we assumed that the diversity of mural cell’s functions depends on their proximity and interaction degree to other vascular cells, which is reflecting cellular collaboration in NVU.

### 4. Chronic METH exposure triggers neuroinflammation through coordinated interactions among immune cells in the central nervous system and peripheral compartments

Traditional METH research has always linked neuroinflammation to astrocytes, microglia, or certain neuroinflammatory factors, but in recent years, the functions of immune cells residing in neural tissue and the neural immune system deserve our attention. Different neural inflammatory factors or immune signals drive different cellular responses through various signaling pathways. As specialized immune cells of the central nervous system, microglia dominate neuroinflammatory processes under chronice METH exposure and play roles as initiators and executors. We found that microglia in our model up-regulates immune and inflammatory responses, such as MHC class II reaction, IL-1 / IL-2 / IL-6 / TNF superfamily cytokines production, and we also noticed that microglia strengthen communications with astrocytes, leukocytes, and lymphocytes whose bioprocesses include activation, differentiation, chemotaxis and migration. It has provided that TNF-α, IL-1α and C1q released by microglia can induce astrocytes to transform into neurotoxic phenotype cells(69). Recent researches pay more attention to the interaction of central and peripheral immune cells influencing neurodegenerations by various mechanisms. Xiaoying Chen et al. reported activate microglia encoding more MHC Ⅰ and Ⅱ proteins recruit T cells into brain parenchyma resulting encephalanalosis(70). Besides, by these innate immune factors, microglia can react to METH-induced DAMPs (Damage Associated Molecular Patterns) from neurons or other cells. We also noticed that some of characteristic cellchat signals induced by METH (such as TIGIT, RANKL, SN, SPP1) are mainly caused by these peripheral immune cells espcially NK-T cells. Astrocytes reshape their morphological, genomic, metabolic, and functional characteristics in response to acute or chronic pathological stimuli through a process called reactive astrogliosis(71). Althought we found astrocytes as a whole can regulate inflammatory response, their functions of different subgroups still need to be classfied by genomic and functional features related to their location and interactions with other cells in the CNS. Specially, we noticed second-messenger-mediated signaling-cAMP is up-regulated in astrocytes, and it may act as a molecular switch for neuroprotective astrocyte reactivity(72). A variety of cells and factors in hippucampal microenvironment participate in or influence neuroinflammation after METH exprosure, our data broadened the research field of neuroinflammation and provided valuable reference genes for various potential inflammatory participants (Fig.9).

**Fig. 9.**
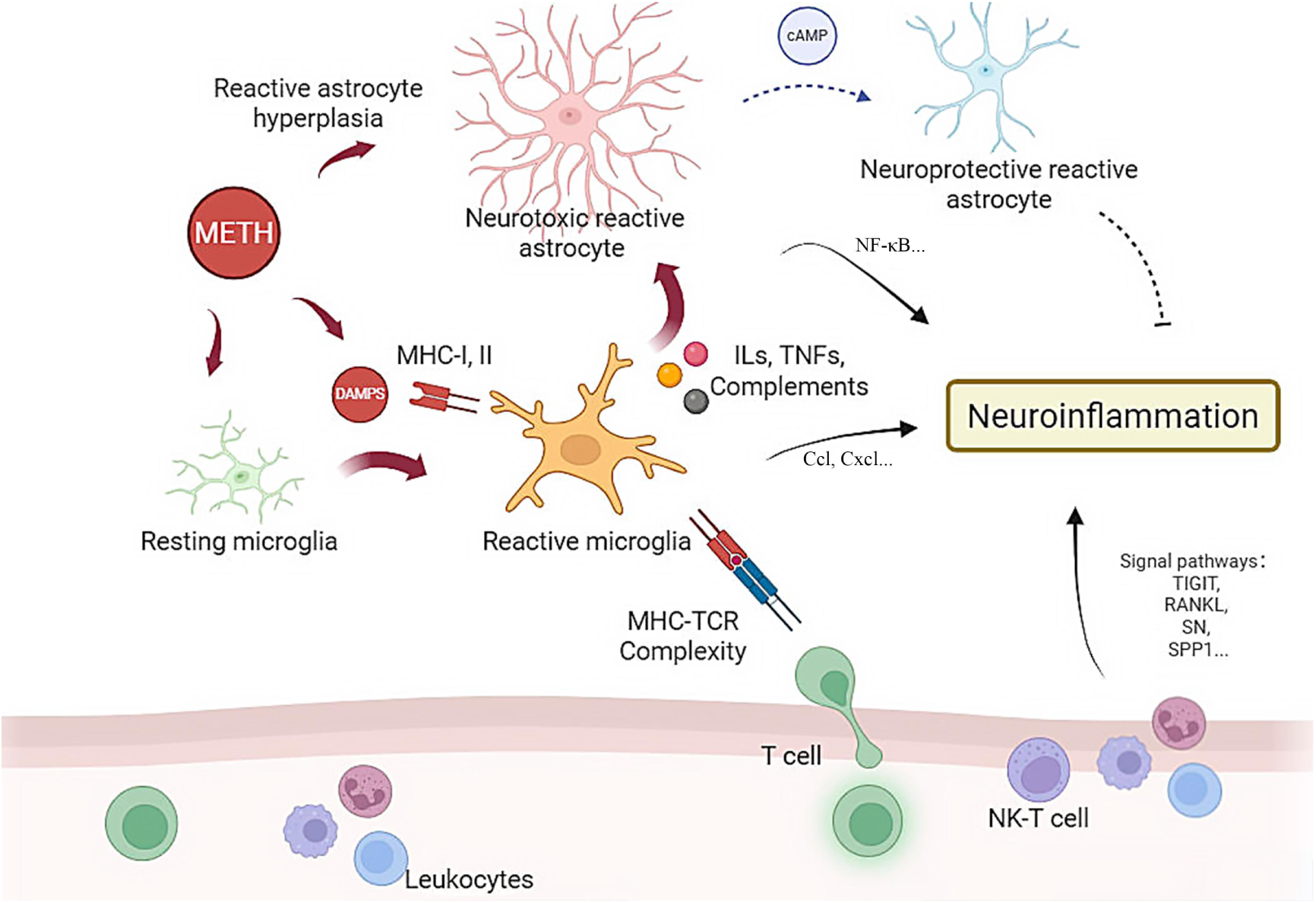
A schematic illustration of the mechanisms underlying neuroinflammation induced by chronic METH exposure. METH induces the reactive activation of microglia and astrocytes via multiple signaling pathways and facilitates the recruitment of peripheral lymphocytes and leukocytes to the central nervous system, thereby contributing to the complex regulatory mechanisms underlying the neuroinflammatory response.

### 5. Identification of disrupted metabolic and circadian coordination across diverse cell types

The metabolism of various cells effected by METH in the hippocampus can not be ignored, as this may also be the cause or result of changes in cell functions. We have paid particular attention to changes in the metabolism of oligodendrocytes, as it is closely related to the support of neural function and structure.we have found that metabolic processes such as the production of organic acids, amino acids, glycerolipids, and glycolipids occur in oligodendrocytes and are similar to the changes in the expression of lipid transport genes emphasized in AD, which may lead to disruptions in myelin formation or energy supply to neurons(73). Furthermore, endothelial cells and Cldn5^+^ enterocytes both showed impairments in lipid transport, localization, and metabolism, involving genes such as *Slc2a1*, *Mfsd2a*, *Spp1*, *Apoe*, the *Atp* family, and the *Abca* family. While current research suggests that the characteristics of the BBB in a healthy state are not dependent on the maintenance of microglial cells, the microglial scavenger PLX5622 has been found to impact endothelial cholesterol metabolism(74). Additionally, impaired lipid metabolism in microglial cells is considered an important pathogenic factor in AD(75). So we hypothesize that endothelial and epithelial cell lipid metabolism and transport disorders triggered by METH may promote lipid accumulation in microglia and cause neurological damage.

The disruption of circadian rhythms has rarely been mentioned in previous studies on METH, but METH, as a neurostimulant drug often used in nightlife, can affect sleep. Interestingly, the frequent occurrence of circadian regulation in our analysis of a number of cell-types is also worthy of attention, e.g. Genes involved in circadian rhythm regulation, like *Per2*, *Per3, Ciart*, *Nr1d1* or *Nr1d2,* are up-regulated in gliocytes such as astrocytes, oligodendrocytes and microglia, *Nrd1*, *Nrd2*, *Sfpq* (regulates the circadian clock by repressing the transcriptional activator activity of the CLOCK-ARNTL heterodimer(76)) are up-regulated in macrophage. Recent research had reported that the circadian clock regulates the immune response of various peripheral innate and adaptive immune cell types and astrocytes and microglia also possess functioning circadian clocks, and circadian timing can affect their inflammatory response, which may be the regulatory mechanism of neuroinflammation in AD(77). Moreover, we noticed that circadian rhythm disorder in endothelia (involving *Per2*, *Cry2*, *Cry1* and *Ptger4*) , smooth muscle cells (involving *Cavin3*, *Kmt2a*, *Per1*, *Nr1d2* et al.) and ependymal cells (involving *Phlpp1*, *Ptgds*, *Clock*, *Id2*). Existing researches showed that the clearance of metabolites, transporter function, permeability, vascular inflammatory and vasomotion of BBB are regulated by circadian rhythm(78–80). A few studies of METH abuse were concerned about circadian rhythm disorder and reported that METH-associated heart failure(81), variability of body temperature(82), Learned motivation(83), addictive properties(84) and learning and memory impairments(28) are affected by disrupted circadian rhythms. However it still needs more researches to figure out the mechanisms of circadian rhythm disturbed by METH and how does circadian rhythm disruption cause METH toxicity.

## Conclusion

In conclusion, our reaserch provides scRNA data on the changes of various types of cells in the hippocampus of mice with cognitive impairment caused by METH exposure. These results provide new insights while partially confirming the results of previous studies on METH neurotoxicity. And we will conduct more confirmatory work in the future to elucidate the mechanisms of neurotoxicity and cognitive decline caused by chronic METH abuse.

## Ethics approval and consent to participate

All protocols approved by the Institutional Animal Care and Use Committee in Southern Medical University (Ethical number: L2022125), and consistent with NIH Guidelines for the Care and Use of Laboratory Animals (8th Edition, U.S. National Research Council, 2011).

## Consent for publication

All authors have approved the manuscript and this submission.

## Authors and Affiliations

School of Forensic Medicine, Southern Medical University, Guangzhou 510515, Guangdong province, People’ s Republicof China: DF Qiao, H Qiu, X Yue, YB Huang, ZL Meng.

Cancer Research Institute, School of Basic Medical Sciences, Southern Medical University, Guangzhou 510515, China: JH Wang.

Corresponding author: Correspondence to JH Wang and DF Qiao.

## Authors’ contributions

H Qiu: Conceptualization, Experiments, Data Collection, Statistical Analysis, Sing cell Data Analysis, Writing (Original Draft), Writing (Editing). X Yue, YB Huang and ZL Meng: Animal treatments and Behavior data analysis. JH Wang: Sing-cell Data Analysis, Funding. DF Qiao: Supervision, Funding, Writing (Editing).

## Availability of data and material

The single-cell RNA-seq data that support findings of this study are available in Gene Expression Omnibus (GEO) under accession numbers GSE252939. Other experimental data and data analysis code can be obtained from the corresponding author at request.

